# ASTEROID, A Validated Annotated Dataset of Analytical Decisions in Graph Theory–Based fMRI

**DOI:** 10.64898/2026.11.06.764858

**Authors:** Sana Hassan Imam, Daniel Kristanto, Daniela Rodríguez De Castro, Inga Henrike Zimmermann, Julius Welzel, Cassie Ann Short, Carsten Gießing, Andrea Hildebrandt

## Abstract

Graph theory-based functional magnetic resonance imaging (fMRI) studies use complex analytical pipelines, but methodological decisions are often reported in unstructured text, limiting systematic comparison and the design of multiverse analyses. Here, we describe ASTEROID, a structured annotation dataset derived from 156 graph theory-based human fMRI study publications. Its schema contains 125 labels, comprising 61 methodological labels and 64 labels for study characteristics and experimental design. An LLM-assisted workflow generated initial annotations, which underwent automated quality-control checks and expert human review. The dataset captures preprocessing, functional connectivity, and graph-analysis decisions. Within these categories, steps and their sequence, software, algorithms, parameters, and supporting text spans are included where available. This accompanying paper provides prompt specifications, processing scripts, and documentation, and describes an interactive viewer that supports exploration of the annotation schema and the study records. ASTEROID enables comparison of reported analytical choices, the informed design of multiverse-based robustness analysis and supports the development and evaluation of automated methodological information-extraction tools and educational resources for fMRI data analysis.

## Background

Graph theory–based analysis of functional magnetic resonance imaging data (fMRI) is a central approach for studying the large-scale organization of functional brain networks. In this framework, brain networks are represented as graphs, in which nodes correspond to brain regions and edges represent statistical dependencies between them^1^. This representation allows researchers to quantify brain network properties such as integration, segregation, modularity, and centrality, which provide insights into functional brain organization and its variation across individuals, cognitive and affective processes, development, and disorders ^2,3^. Graph-theoretic approaches are at the core of network neuroscience, supporting both descriptive accounts of brain organization and predictive investigations of cognition, emotion, and behaviour ^4^.

Functional brain networks cannot be observed directly from raw data; rather, they are constructed through a sequence of data processing and analysis decisions, commonly referred to as an analytical pipeline. In graph theory-based fMRI analysis, an analytical pipeline typically comprises multiple preprocessing steps of the brain images, including, but not limited to, motion correction, normalization, and filtering ^5–7^, followed by graph analysis steps, such as connectivity estimation, thresholding, and graph-metric computation ^8–10^. Most steps offer multiple defensible options, each of which can alter the data and the resulting network representation. For example, motion correction and denoising choices and their modular workflows can alter residual artifacts in the data. Similarly, choices of connectivity measures and network construction methods can substantially alter the estimated network structure ^8–10^. Consequently, variations in preprocessing and graph analysis steps can lead to substantially different results.

The numerous steps involved in graph theory-based fMRI analysis pipelines, together with the multitude of defensible options available at each step, create a broad space of plausible analytical pipelines that was termed the ‘garden of forking paths’ ^11^. In the traditional one-pipeline analysis approach, researchers select a single workflow from this decision space, leaving considerable ‘researcher degrees of freedom’ in how raw data are transformed into final reported results uncovered ^12^. This flexibility is evident in the graph theory-based fMRI literature, where analytical pipelines vary considerably in the inclusion, ordering, and parameterization of analytical steps^13^. A large, multi-analyst neuroimaging study demonstrated the practical consequences of such flexibility, wherein 70 independent teams analysed the same fMRI dataset to test the same hypotheses, yet no two teams used identical pipelines, and their reported results varied substantially^14^. These findings suggest that low replicability in neuroimaging may be partly due to heterogeneity in how data are processed and analysed. Therefore, the facilitation of robustness analysis in this field requires an explicit, structured overview of the analytical decision space.

Multiverse analysis makes the uncertainty of analytical flexibility transparent by applying all, or a principled subset of all, analytical pipelines to the same dataset and reporting the robustness of the results across all computed pipelines^15,16^. However, conducting a multiverse analysis requires researchers to first map the garden of forking paths for a given dataset and research question^17^. Published pipelines can provide empirical guidance for defining this decision space^13^, but analytical decisions are typically reported in unstandardized prose, described using inconsistent terminology, and reported with varying levels of detail ^18,19^. Standards such as the Brain Imaging Data Structure (BIDS) have improved the organization and sharing of neuroimaging datasets and associated metadata ^20^, however they do not resolve the substantial researcher degrees of freedom. Recent work further demonstrates that analytical variability also extends to AI-based analysts, which can produce divergent analytical choices and outcomes when addressing the same research question^21^.Therefore, a structured representation of published analytical pipelines is needed to make the existing analytical decision space searchable, comparable, and reusable ^22^.

Our previous systematic review provided an essential first step toward making analytical flexibility in graph theory-based fMRI explicit ^13^. By mapping preprocessing and graph analytical choices across the literature, the review identified 61 distinct analytical steps and 102 parameter settings, and substantial heterogeneity in the inclusion or omission of specific preprocessing steps, parameterization choices within steps, and the ordering of steps. This manually curated information was integrated into the METEOR decision-support tool to facilitate informed multiverse analyses. However, that resource was designed primarily as a taxonomic synthesis of analytical variability and as a basis for decision support. It did not provide a dataset in which each individual study is annotated using a standardized schema, nor did it link each reported analytical decision to the exact sentence or paragraph in which it was described in a given paper. Such a dataset is needed to enable systematic comparison of published pipelines, transparent verification of annotations, and future benchmarking of automated information extraction on methods from the published literature.

Here, we describe ASTEROID, an **A**nnotated and **S**tructured da**T**as**E**t of g**R**aph-based fMRI analytical **O**pt**I**ons and **D**ecisions. The dataset comprises 156 studies and represents methodological information using a structured schema of 125 labels describing analytic decisions and study-context metadata. Building on advances in natural language processing (NLP) and large language model (LLM)-based information extraction from scientific texts^23,24^, we used an LLM-assisted, human-in-the-loop workflow to transform the manually curated literature knowledge space^13^ into machine-readable and validated annotations. The resulting dataset converts heterogeneous narrative descriptions into a standardized, structured dataset for systematic evaluation.

Researchers can use ASTEROID to identify which preprocessing and graph-analysis choices are commonly reported across studies, examine how analytical decisions vary across study designs or populations, and select empirically grounded options when defining the decision space for multiverse analyses. Because annotations are linked to supporting text spans, users can verify how specific decisions were reported in the source literature rather than relying only on summarized labels. ASTEROID can also serve as a benchmark for developing and evaluating automated methods that extract methodological information from scientific articles, and as a teaching resource for illustrating how analytical pipelines are reported, structured, and compared across studies. To facilitate access and reuse, the dataset is accompanied by an interactive exploration interface that allows users to inspect the annotation schema, explore annotated studies, and compare pipelines across the literature. Together, the dataset and interface provide a resource for studying analytical heterogeneity and changes in research practice over time. They support transparent and systematic multiverse analyses, while also facilitating education and training. By making published graph theory–based fMRI pipelines more searchable, comparable, and reusable, we improve access to methodological knowledge in the field.

In this data descriptor, we introduce the ASTEROID dataset and provide the information needed to evaluate and use it. We first define the annotation schema and the corpus from which the dataset was derived. We then report the LLM-assisted, human-in-the-loop workflow used to generate and validate the annotations, including the automated quality-control procedures and expert review steps used to ensure consistency. Finally, we describe the data records and the interactive interface. We show how the resource can be used to inspect annotation structures, trace analytical decisions to supporting text, compare pipelines across studies, and assess the dataset’s quality and scope. By documenting both the dataset and its validation, this data descriptor increases the transparency and usability of the dataset.

## Methods

The dataset creation process comprised five main stages: corpus collection, PDF-to-JSON conversion, schema-guided LLM-assisted pre-annotation, automated quality control, and consensus-based expert validation. This workflow was designed to transform narrative methodological descriptions from published articles into a structured JSON dataset with standardized labels, attributes, and supporting text spans. The full dataset construction workflow is illustrated in Figure 1, and each stage is described in detail below.

**Figure 1:**
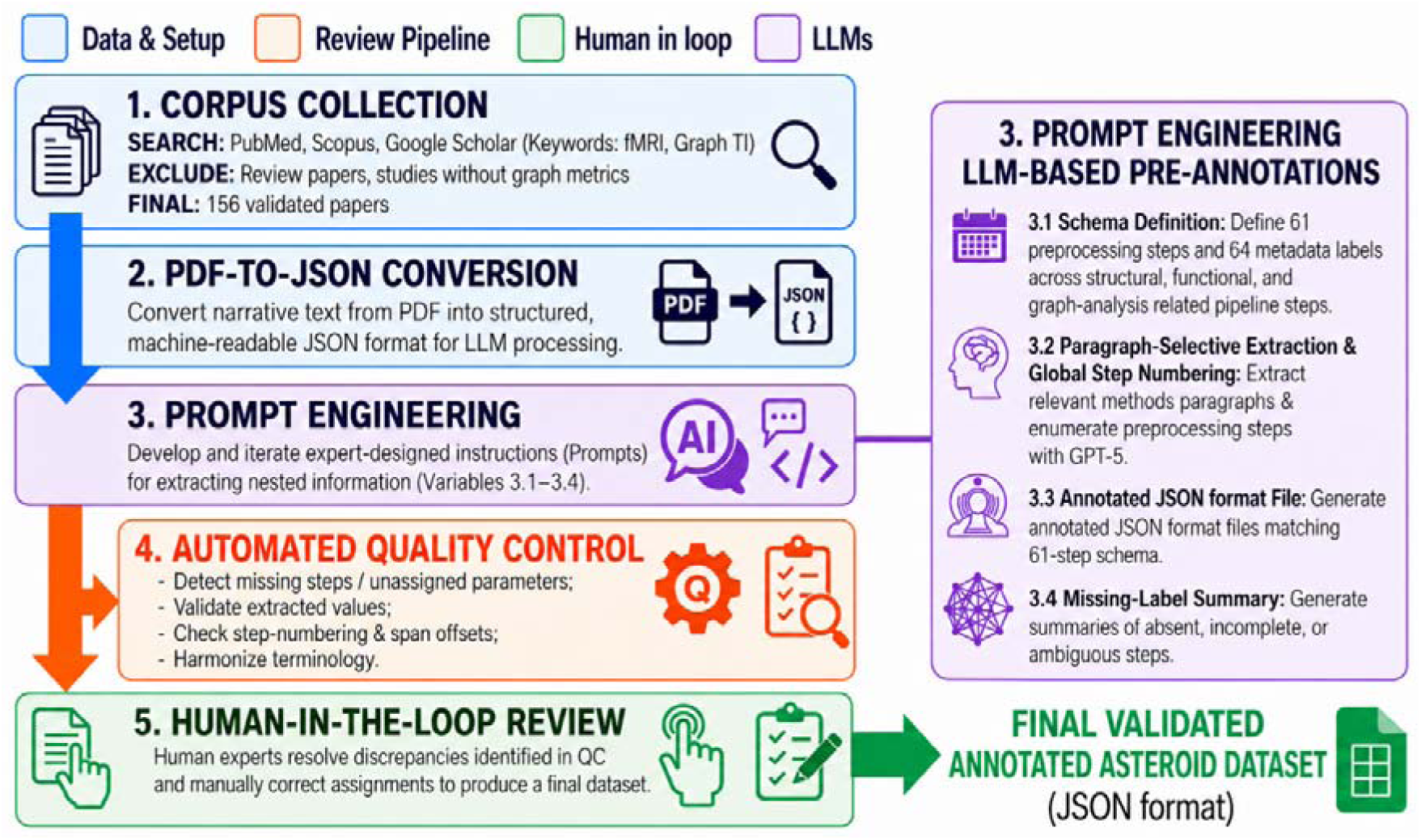
Workflow for constructing the ASTEROID benchmark dataset. Overview of the multi-stage workflow used to create the validated and annotated dataset of graph theory–based fMRI analytical decisions. The process includes corpus collection, PDF-to-JSON conversion, LLM-assisted pre-annotation using a predefined schema, automated quality control, and expert human review. The final output is the validated ASTEROID dataset. Colors indicate workflow stages: data & setup (blue), review pipeline (orange), LLM-assisted (purple), and human in loop (green).

### Corpus collection

The graph-theory-based fMRI studies to be analysed were collected from PubMed, Scopus, and Web of Science^13^. All collected documents were stored in a directory as PDFs, forming the raw corpus for subsequent processing.

### PDF-to-JSON conversion

All full-text PDFs were converted into structured JSON files using a standardized parsing script to support paragraph-level LLM-assisted extraction. The conversion script is available at <u>Github: Annotation-buddy</u>. The resulting JSON files contained the extracted text and document-level information, with automatic removal of headers, footers, and repeated section labels. Conversion quality was checked by inspecting a subset of converted files for preservation of section headings, paragraph order, and relevant methods text. The annotation process focused primarily on main-text methods sections and related analysis descriptions. Tables, figures, captions, reference lists, and supplementary materials were not systematically annotated, but were consulted when they contained information needed to clarify reported methodological steps. This approach ensured that the final annotations were mainly grounded in the narrative methodological descriptions reported in the published articles.

### Prompt engineering and LLM-assisted pre-annotation

Each full-text JSON was processed with a prompt applied to GPT-5 through the ChatGPT interface (OpenAI), accessed between August 2025 and February 2026. The prompt implemented the following: (i) schema definition, (ii) paragraph-selective extraction of methodological content, (iii) mapping of preprocessing steps to a standardized schema with generation of an annotated JSON file, and (iv) creation of a missing-label summary for quality control. The prompt has been refined iteratively to improve the quality of extraction of targeted study-characteristics information, including publication details, participant characteristics, data acquisition parameters, task and experimental design features, and analytical pipeline decisions. The LLM-generated outputs were treated as pre-annotations and subjected to automated quality control and expert human review before inclusion in the final dataset. The prompt versions are provided in the Prompts/ directory of the ASTEROID GitHub repository, described in the Data Records section.

#### i. Schema definition

In response to the lack of standardization in reporting analysis pipelines in the fMRI literature, we developed a multi-layered annotation scheme comprising 125 distinct labels. Figure 2 summarizes the main label groups and selected example labels rather than displaying the full set of 125 labels. The scheme is organized from left to right into three main components: **Study characteristics**, **Experimental design**, and **Analytical pipeline**.

**Figure 2.**
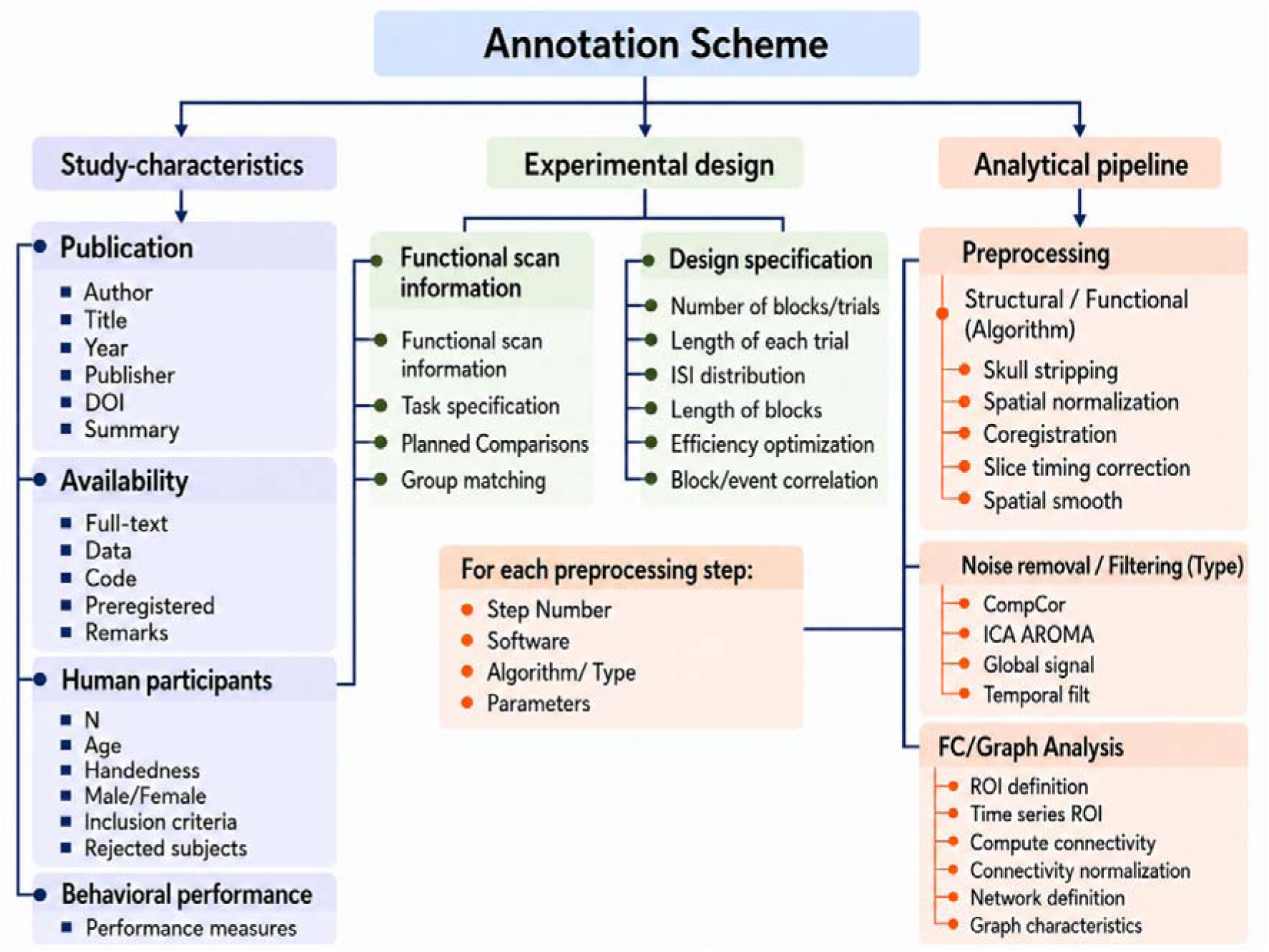
Hierarchical annotation schema for graph theory–based fMRI analytical decisions. The schema contains 125 labels organized into three major components: study-characteristics, experimental design, and analytical pipeline information. The figure should be read from left to right, moving from general study-characteristics information to the reported analytical workflow. For each preprocessing or analysis step, the schema records four standardized attributes: step number, software, algorithm/type, and parameters.

The first component, **Study characteristics**, captures publication details, availability information, human-participant characteristics, and behavioral performance measures. The second component, **Experimental design**, captures functional scan information and design specifications, including task-related information and trial/block-level details. Together, **Study characteristics** and **Experimental design** comprise 64 study-characteristic labels, which provide the contextual information needed to interpret the methodological choices reported in each study.

The third component, **Analytical pipeline**, captures 61 methodological labels describing the reported image-processing and analysis workflow. Within this component, Figure 2 illustrates the **Preprocessing** label group, which is further organized into structural/functional algorithm information, noise removal/filtering procedures, and FC/graph-analysis steps. The figure shows representative examples within these groups, such as skull stripping, spatial normalization, CompCor, ICA-AROMA, ROI definition, connectivity computation, network definition, and graph characteristics; however, not all 61 methodological labels are shown in the figure.

For each reported preprocessing or analysis step, four standardized sub-attributes were extracted: step number, software, algorithm/type, and parameters. By linking the **Analytical pipeline** labels with the contextual labels from **Study characteristics** and **Experimental design**, the annotation scheme transforms unstructured methodological descriptions into a harmonized, structured, machine-readable dataset suitable for large-scale comparison, automated information extraction, and meta-scientific analysis.

This uniform attribute structure preserves not only which steps were reported, but also how they were implemented and in what order. It therefore supports reconstruction of complete analytical pipelines and comparison of methodological choices across studies.

#### ii. Paragraph-selective extraction and global step numbering

Paragraph selection was performed by GPT-5 based on prompt instructions. The model received the full article text and selected paragraphs containing methodological information relevant to the schema. When overlapping information occurred across sections, the model prioritized content in the order: Methods → Results → Discussion → Abstract/Introduction. Step numbering followed a single global sequence inferred from linguistic cues and common neuroimaging workflows^13^.

#### iii. Annotated JSON file

The prompt produced an annotated JSON file in which the LLM mapped relevant information from the three schema components, Study characteristics, Experimental design, and Analytical pipeline, to the predefined annotation scheme. Relevant text spans were detected and assigned to the corresponding labels, resulting in a structured, machine-readable output that could be directly imported into HumanSignal and compared with manually extracted annotations from the previous study^13^.

#### iv. Missing-label summary

For each paper, the prompt generated a missing-label summary to identify reporting gaps in the narrative literature. These summaries listed labels from the predefined annotation scheme that were not detected in the paper text. The missing labels were organized according to the three main schema components shown in Figure 2: Study characteristics, Experimental design, and Analytical pipeline. Within the Analytical pipeline component, missing preprocessing-related labels were further grouped under structural/functional algorithm information, noise removal/filtering, and FC/graph-analysis steps.

##### Absent

Targeted information was marked as absent when it was not reported in the text. For example, if a paper did not mention slice timing correction, the corresponding preprocessing label was marked as absent.

##### Incomplete

Targeted information was marked as incomplete when a preprocessing or analysis step was mentioned in the text, but one or more required sub-attributes were missing. For example, if a paper reported spatial smoothing but did not provide the FWHM parameter, the corresponding label was marked as incomplete.

##### Ambiguous

Targeted information was marked as ambiguous when it was reported using non-standard or vague terminology that prevented a definitive mapping to the predefined annotation scheme. For example, if a paper described “standard denoising” without specifying whether this referred to ICA-AROMA, CompCor, or global signal regression within the noise removal/filtering group, the corresponding label was marked as ambiguous.

This process allowed us to systematically track reporting quality across the 156 papers. The missing label summary file supported automated checks and guided later human review.

### Prompt refinement

From the initial literature corpus, 156 human fMRI studies were included in the ASTEROID dataset. During early validation, we observed that core preprocessing steps, such as motion correction, spatial normalization, and smoothing, were usually reported more explicitly than graph-analysis related information. In contrast, details related to atlas/parcellation choice, connectivity metrics, edge definitions, thresholding rules, and graph-construction parameters were often sparse, inconsistently defined, or reported only implicitly.

To address this issue, we used a staged prompt-refinement strategy with three prompt versions: the First Prompt, the Graph Analysis Prompt, and the Final Prompt. The First Prompt was designed as a broad extraction prompt covering study-characteristics information, experimental design details, and analytical pipeline decisions. It was applied to the first 80 studies. Review of these outputs showed that graph-analysis information required more explicit domain-specific guidance. We therefore developed the additional Graph Analysis Prompt, which focused on graph-analysis steps and included more detailed instructions for extracting information on connectivity computation, network construction, thresholding, graph measures, and related parameters.

Outputs from the first prompt and the graph analysis prompt were merged using a custom Python script to improve coverage of graph-analysis information while retaining the broader study-characteristics labels and preprocessing annotations. For the remaining 76 studies, we used the final prompt, which incorporated the improvements from the earlier prompt versions and covered the full 125-label schema in a single pass. This staged approach supported more consistent extraction across heterogeneous reporting practices in the source literature, while all outputs remained subject to automated quality control and expert human review before inclusion in the final dataset.

### Review Pipeline

The review process for the LLM-generated annotations followed a two-stage "human-in-the-loop" pipeline to ensure the annotations’ consistency and accuracy (see Figure 3).

**Figure 3:**
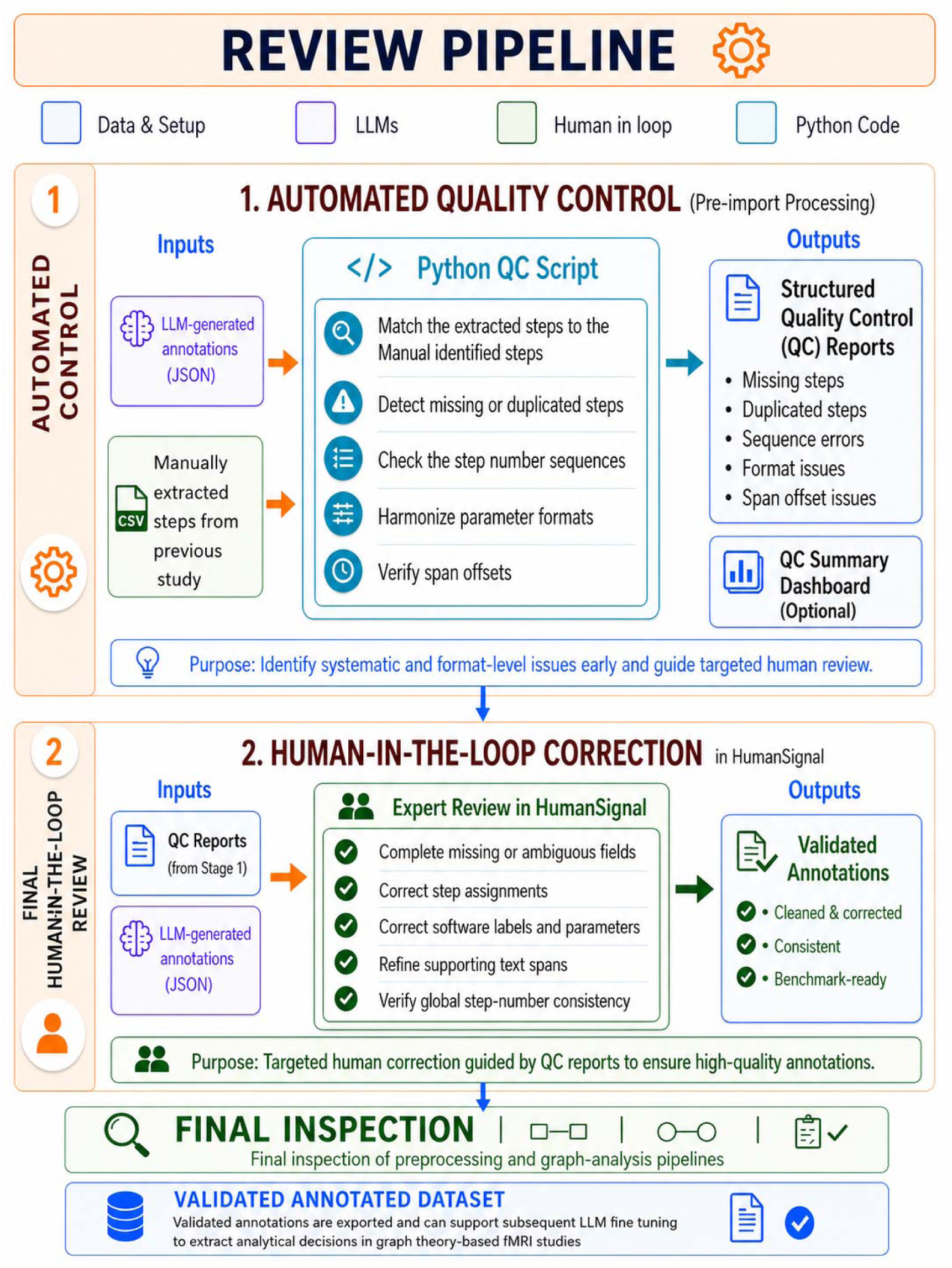
Human-in-the-loop review and validation workflow. Automated quality-control (QC) scripts compared pre-annotations with manually curated reference information, detected missing, duplicated, inconsistent, or ambiguous entries, and generated structured review reports. Expert reviewers then inspected and corrected the annotations in HumanSignal/Label Studio, including label assignments, step attributes, parameters, spans, and global step numbering. The reviewed annotations were exported as the final validated dataset ASTEROID.

#### Automated quality control

Before importing the annotated JSON files into HumanSignal, a Python script was applied to the LLM-generated annotations. The script matched extracted steps to manually identified steps from our previous study^13^, detected missing or duplicated steps, checked step-number sequences, harmonized parameter formats, and verified span offsets. All discrepancies were compiled into structured quality control reports.

#### Final human review and validated dataset assembly

In the final round, as shown in Figure 3, the annotated JSON files were imported into HumanSignal for expert review. Guided by quality control reports, the first and third authors and a student assistant reviewed, corrected, and finalized the annotated files by completing missing or ambiguous fields, correcting step assignments, software labels, and parameters, refining supporting spans, and verifying global step-number consistency. After this targeted human-in-the-loop correction, a final inspection of the preprocessing and graph-analysis pipelines was performed. The review was not designed as an independent parallel annotation task; therefore, inter-annotator agreement was not calculated. Uncertain cases were resolved through discussion among the reviewing team. All other co-authors additionally reviewed the final structural workflow of the annotation process to assess its clarity, completeness, and consistency. The validated annotations were exported as JSON files, forming the validated annotation dataset, which can support subsequent LLM fine-tuning to extract analytical decisions from graph theory-based fMRI studies.

## Data Records

The final ASTEROID dataset contains validated annotations from 156 graph theory–based human fMRI studies. The ASTEROID dataset and accompanying documentation have been deposited on Zenodo under the reserved DOI (https://doi.org/10.5281/zenodo.23569927). The dataset is currently under review and will be made publicly available upon completion of the necessary release requirements. The repository will provide the validated annotations, schema documentation, processing scripts, and supporting materials required for reuse and interpretation. The associated project repository is hosted on GitHub (https://github.com/sanahassanimam/ASTEROID-Validated-Annotated-Dataset) and will be made publicly accessible upon dataset release.

The main data file is stored as:

Annotated_Files/all_annotations.json

This file contains one structured record per study and is compatible with the Label Studio JSON import format. Each record includes **Study characteristics**, **Experimental design**, and **Analytical pipeline** annotations, along with step-specific attributes and supporting text spans where available. The unified JSON format allows users to inspect individual studies, compare reported pipelines across studies, and import the data into compatible annotation or analysis platforms.

The repository is organized to support transparent reuse of the dataset and its supporting materials (Figure 4). It includes the validated annotations, prompt templates, intermediate LLM-generated outputs, Python scripts for merging and quality control, figure resources, and an interactive schema viewer.

**Figure 4.**
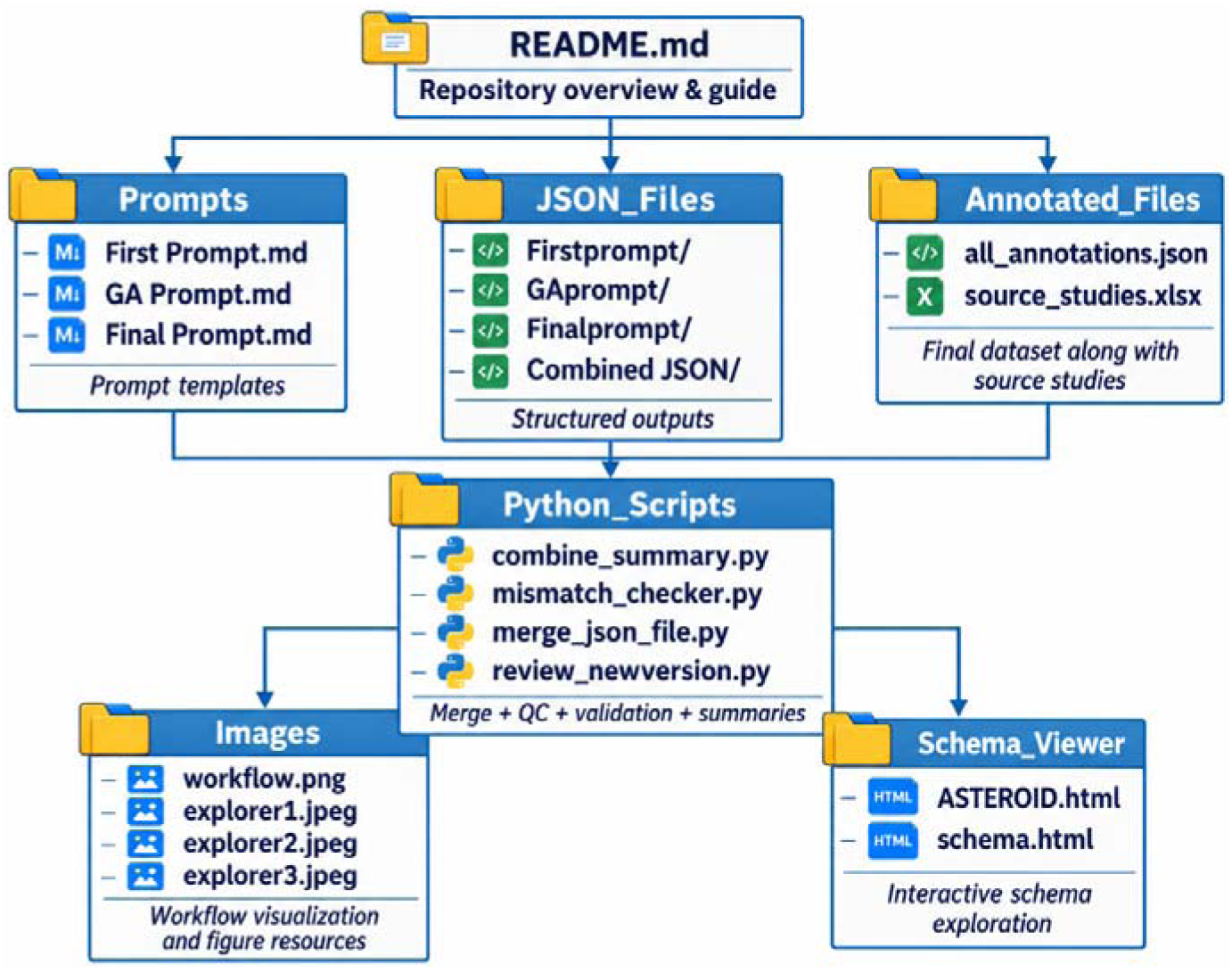
Repository architecture and dataset organization. The repository contains the final validated annotations, prompt templates, intermediate LLM-generated JSON outputs, Python scripts, figure resources, and an interactive schema viewer. The main dataset is stored in Annotated_Files/all_annotations.json. Supporting folders document the pre-annotation, merging, validation, and visualization resources used during dataset construction.

The Annotated_Files/ folder contains the final human-validated annotations. The JSON_Files/ folder contains intermediate LLM-generated outputs from different prompt versions, including first prompt, graph analysis (GA) prompt, final prompt, and combined outputs. These files document the pre-annotation process and allow comparison across prompt versions.

The Prompts/ folder contains the prompt versions used during annotation development. These include the first prompt, the GA prompt, and final prompt. They are provided to document how pre-annotations were generated and to support future adaptation of the extraction workflow.

The Python_Scripts/ folder contains scripts used to merge JSON outputs, generate summaries, compare annotations with the manually curated reference database, identify mismatches, and support review of updated extraction results. These scripts document the main processing and validation steps used to prepare the final dataset.

The Schema_Viewer/ folder includes ASTEROID.html and schema.html, which provide browser-based interfaces for exploring the annotated dataset and its underlying annotation schema. The viewer allows users to inspect the annotation schema, explore annotated records and highlighted spans, and generate structured summaries of preprocessing and graph-analysis pipelines. It runs on the client side and does not require backend setup.

The Images/ folder contains workflow diagrams and visualization resources used to illustrate the dataset construction process and the interactive schema exploration interface. These materials provide visual documentation of the annotation workflow and supporting tools.

Together, the repository and annotation schema define the structure of ASTEROID. The dataset provides validated study-characteristics annotations, while the supporting files document the annotation workflow, validation procedures, and tools for exploring the annotated records.

## Technical Validation

Technical validation was organized around two main components. First, we assessed the alignment of the LLM-generated annotations with an independently compiled manual extraction resource. This analysis evaluated annotation coverage, agreement, and the effect of prompt refinement. Second, we assessed the lexical consistency of the extracted text snippets within individual schema labels. This analysis examined whether annotations assigned to the same label captured comparable methodological content across studies. Together, these analyses evaluated both the external agreement and the internal coherence of the ASTEROID dataset. Automated structural checks and expert review of the source articles supported them. The validation procedure was used not only to quantify extraction performance, but also to identify omissions, conflicting values, duplicate records, inconsistent schema use, and labels associated with substantial variation in methodological reporting.

Before the two main validation analyses, all extracted JSON files underwent automated quality control using custom Python scripts. The scripts verified that preprocessing and analysis steps followed a consistent numbering order within each article and across the corresponding structured records. We also detected entries, duplicated annotations, and semantically overlapping records, including cases in which the same methodological step had been extracted more than once under slightly different labels or wording. Parameter formats and naming conventions were harmonized, and annotation offsets were checked to confirm that each extracted record remained aligned with its supporting text span.

All flagged cases were subsequently reviewed by domain experts in HumanSignal/Label Studio. Reviewers compared each extracted span with the original article text, corrected label assignments, resolved mismatches, removed false-positive annotations, and completed missing fields when the source article supported the information. Records found only in the LLM-generated output were examined to determine whether they represented valid additional methodological information, while records found only in the manual extraction were reviewed to distinguish genuine omissions from differences in interpretation or schema structure.

### Alignment with manual database

The first component of the technical validation assessed the alignment between the LLM-generated JSON annotations and the methodological information manually extracted in our previous study^13^. The manually curated database was used as an independent comparison standard rather than as absolute ground truth. This distinction was important because disagreements could reflect omissions, differences in interpretation, or differences in schema structure in either resource. All unresolved cases were therefore adjudicated against the original article text.

Records were classified as Match, Not match, Manual only, or LLM only. Match indicated that the LLM-generated output and the manually curated database contained equivalent information for the same extraction field or methodological decision. Not match indicated that comparable records contained conflicting, incomplete, or semantically non-equivalent values. Manual-only records referred to information present in the manually curated database but absent from the LLM-generated output. LLM-only records referred to information extracted by the LLM but not represented in the previous manual database. LLM-only records were not treated automatically as false positives; each was reviewed against the source article to determine whether it represented valid additional information.

These comparisons formed the basis of the quantitative validation shown in Figure 5 and were also used to evaluate the effect of prompt refinement.

**Figure 5.**
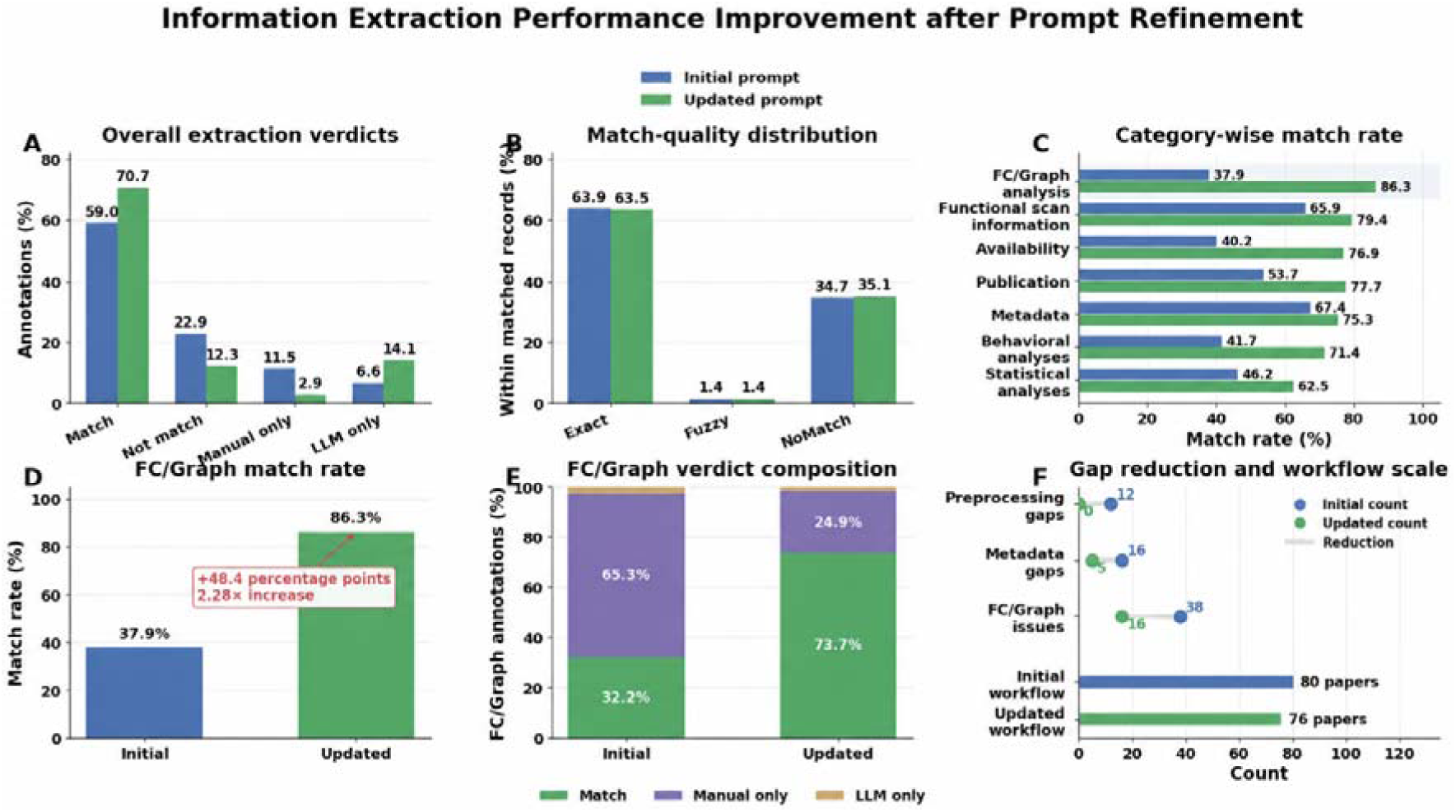
Quantitative comparison of LLM-generated JSON annotations against the manually curated reference database before and after prompt refinement. The manually curated reference database was used as a comparison standard to evaluate the agreement, completeness, and consistency of the LLM-generated annotations. (A) Overall verdict distribution across extracted annotations, where Match indicates annotations present in both the LLM-generated JSON output and the manually curated reference database, Not match indicates comparable annotations with conflicting or non-equivalent values, Manual only indicates annotations present only in the reference database, and LLM only indicates annotations extracted only by the LLM. (B) Record-level match-quality distribution for comparable annotation pairs, distinguishing exact matches, fuzzy/semantically equivalent matches, and non-matching paired records. (C) Category-wise comparison of annotation match rates. (D) Focused comparison of functional connectivity/graph-analysis extraction performance. (E) Composition of functional connectivity/graph-analysis verdict categories. (F) Workflow-level reduction in unresolved preprocessing, study-characteristics descriptor, and functional connectivity/graph-analysis gaps, together with the number of papers processed under each workflow.

#### Effect of prompt refinement

Prompt refinement was evaluated within the manual-alignment analysis because the independently compiled reference database provided a direct basis for assessing whether changes to the extraction procedure improved annotation completeness. For the first 80 studies, outputs from the baseline prompt and a separate graph-analysis-focused prompt were merged into a single structured JSON file for each study. This two-stage workflow increased the coverage of graph-analysis information but still left unresolved discrepancies across preprocessing, study characteristics, and functional-connectivity or graph-analysis categories. For the remaining 76 studies, a refined single-pass prompt was used. This prompt incorporated explicit definitions of functional-connectivity and graph-analysis concepts into the main extraction workflow and imposed clearer constraints on the expected content and structure of the extracted records. The refined prompt was designed to reduce omissions and inconsistencies while removing the need to merge outputs from separate extraction passes.

Before final human correction, the updated prompt increased the overall proportion of matched annotations from 59.0% to 70.7% and reduced the proportion of non-matching annotations from 22.9% to 12.3% (Figure. 5A). Manual-only records decreased from 11.5% to 2.9%, indicating that the refined prompt recovered a substantially larger proportion of the methodological information represented in the comparison database including manually extracted information. The proportion of LLM-only records increased from 6.6% to 14.1%. Because the reference database was not treated as absolute ground truth, these records were examined individually to determine whether they represented false-positive extractions or valid methodological details that had not been captured during the earlier manual review.

Among records that could be aligned across the two resources, the distribution of match quality remained broadly stable across prompt versions (Figure. 5B). Exact matches accounted for 63.9% of comparable pairs with the initial prompt and 63.5% with the updated prompt. Fuzzy or semantically equivalent matches accounted for 1.4% in both workflows, while non-matching comparable pairs accounted for 34.7% and 35.1%, respectively. In this analysis, NoMatch refers to record pairs that could be aligned for evaluation but whose extracted values did not agree. It does not refer to a subset of otherwise matched annotations.

The stability of the match-quality distribution indicates that prompt refinement primarily improved annotation coverage by recovering information that had previously been missed. It did not substantially alter the agreement profile of records that could already be paired across the two resources.

The largest improvement was observed for functional-connectivity and graph-analysis annotations. Their match rate increased from 37.9% with the initial prompt to 86.3% with the updated prompt, corresponding to a 48.4-percentage-point increase and a 2.28-fold relative improvement (Figure. 5C, D). The composition of functional-connectivity and graph-analysis verdicts changed accordingly: matched records increased from 32.2% to 73.7%, while manual-only records decreased from 65.3% to 24.9% (Figure. 5E). These results show that explicitly defining functional-connectivity and graph-analysis concepts within the extraction prompt substantially improved the recovery of complex downstream analytical decisions.

The effect of prompt refinement was also evident at the workflow level. Unresolved preprocessing gaps decreased from 12 to 0, metadata-related gaps decreased from 16 to 5, and functional-connectivity or graph-analysis issues decreased from 38 to 16 (Figure. 5F). These reductions were observed even though the updated workflow was applied to 76 papers, compared with 80 papers processed using the initial workflow. The refined prompt therefore reduced the number of unresolved issues per study and decreased the amount of manual correction required during final validation.

The manual-alignment analysis shows that prompt refinement improved both the completeness of the automated extraction and its agreement with the independently compiled reference resource. The improvement was particularly marked for graph-analysis information, which is often distributed across different sections of an article and reported using variable terminology. Remaining disagreements were resolved through article-level expert review so that the final annotations reflected the source publications rather than either extraction resource alone.

Together, these results show that automated quality control, prompt refinement, and expert human-in-the-loop review improved the completeness, agreement, and internal consistency of the final ASTEROID annotations. The validation process was therefore used not only to assess extraction performance against the manually curated reference database, but also to guide correction, harmonization, and consolidation of the final ASTEROID dataset.

### Lexical consistency of extracted snippets

To assess whether text snippets assigned to the same schema label describe comparable methodological content, we quantified the lexical consistency of the snippets within each label. Whereas the comparison with the manually curated database evaluated whether the extraction workflow recovered previously identified methodological information, this analysis assessed the internal consistency of annotations grouped under the same schema label.

Across the 156 annotated studies, this yielded 3,625 text snippets distributed across 55 methodological labels (a total of 61 methodological labels are mentioned above, of which 6 were not reported in any study). All snippets were represented as term-frequency–inverse-document-frequency (TF-IDF)^25^ vectors in a single shared vector space fitted over all analysed snippets, ensuring that cosine-similarity scores were directly comparable across labels. For every label represented by at least five snippets, we computed the cosine similarity of all unique snippet pairs within that family, yielding 289,143 pairwise comparisons in total. We summarized each label family using the median cosine similarity of its term-sharing pairs, defined as pairs sharing at least one vocabulary term and therefore having a cosine similarity greater than zero. The proportion of non-sharing pairs was reported separately. This distinction prevented pairs with complete lexical non-overlap from dominating the summary statistic and allowed variation among snippets sharing relevant terminology to be examined more directly.

The ranking of the most and least consistent labels was restricted to label families containing at least 30 snippets because, for small families, a limited number of similar snippets could disproportionately influence the statistic. At a threshold of five snippets, the ranking was significantly associated with the number of snippets per family (Spearman’s ρ = −0.33, *p* = 0.037). At a threshold of 30 snippets, this association was no longer significant (ρ = +0.18, *p* = 0.367), supporting the use of this threshold for comparative ranking.

We evaluated the lexical consistency of the schema labels based on the distribution of within-label cosine similarities (Figure. 6). The results revealed a clear methodological division between standardized structural and functional preprocessing procedures and more heterogeneous graph-analysis decision descriptions. Segmentation, skull stripping, and slice-timing correction were the labels most consistently described in lexical terms, with median cosine similarities above approximately 0.4. Researchers commonly perform these tasks using established algorithms and standard software packages, resulting in relatively consistent methodological terminology and reporting. Their higher within-label similarity therefore supports the coherence of these annotation classes and indicates that snippets assigned to them generally describe comparable analytical operations.

**Figure 6.**
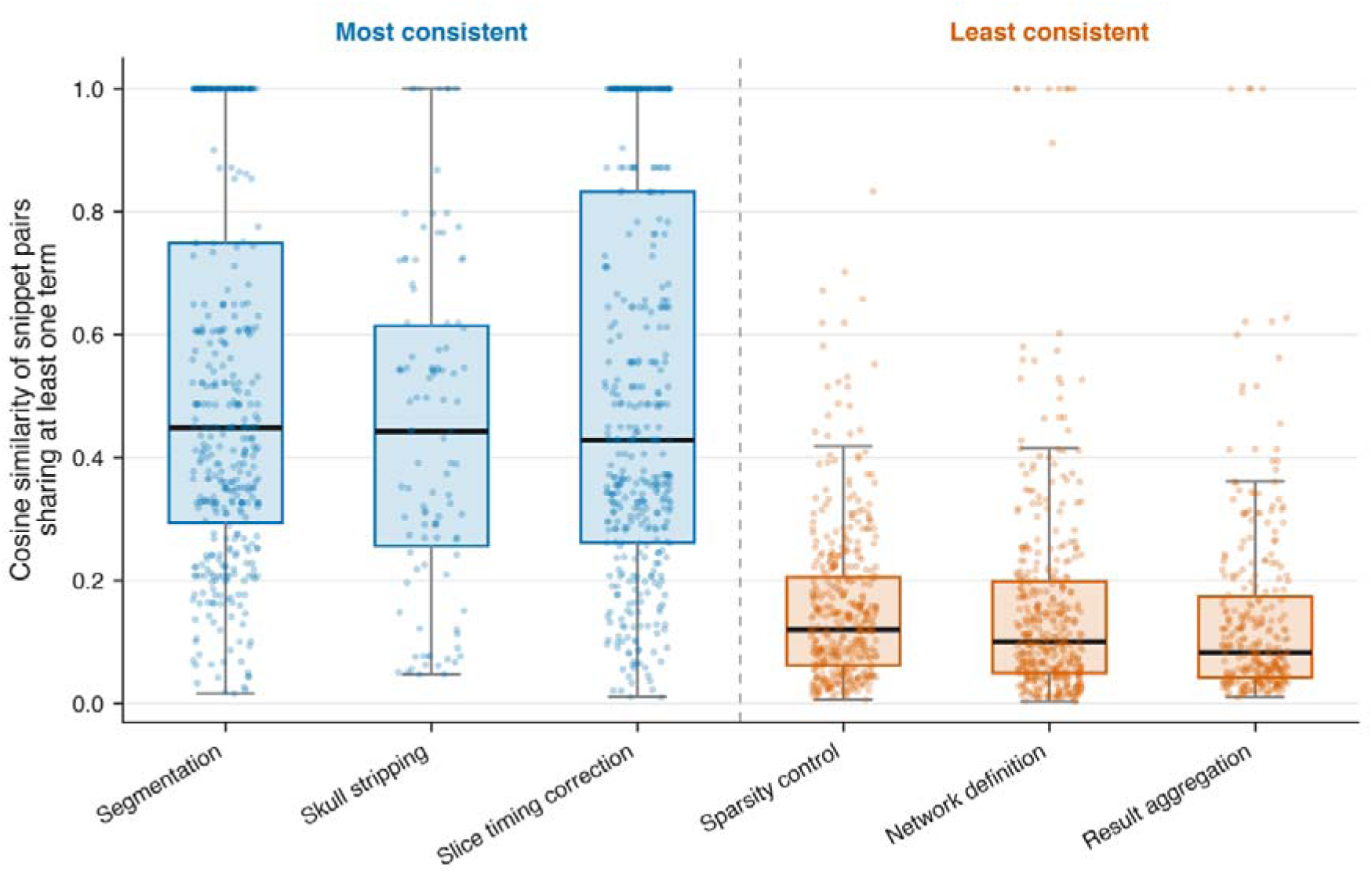
Within-Label Similarity of Term-Sharing Annotated Text Snippet Pairs for the Most and Least Consistent Labels. Pairwise cosine-similarity distributions for term-sharing snippets assigned to the same schema label. Segmentation, skull stripping, and slice-timing correction showed the highest median similarities and represent comparatively standardized preprocessing procedures. Sparsity control, network definition, and result aggregation showed the lowest median similarities and represent graph-analysis decisions characterized by greater methodological and terminological variation. Points represent individual snippet pairs, boxes indicate the interquartile range, and the central line indicates the median.

By contrast, the least lexically consistent labels belonged to the graph-analysis phase. Sparsity control, network definition, and result aggregation showed median cosine similarities of approximately 0.1. Unlike standardized structural and functional preprocessing steps, these stages offer greater flexibility in their implementation, parameterization, and description. We defined networks using different anatomical or functional criteria, applied different sparsity thresholds or thresholding strategies, and aggregated graph-analysis results using study-specific reporting structures. To justify these methodological choices, we used variable narrative structures and custom terminology. Consequently, snippets assigned to these labels may describe methodologically related decisions while sharing relatively little surface vocabulary.

The lower lexical similarity of these graph-analysis labels should not, by itself, be interpreted as evidence of poor annotation quality. Instead, it indicates that the same methodological concepts are reported using more varied language across the literature. These text-based findings are supported by the empirical results of our previous systematic review^13^, in which we identified these graph-theoretical stages as involving multiple and sometimes debated parameter choices. These findings can also be explored in the METEOR app by navigating to the “Steps: Parameter” tabs and selecting the steps with the lowest within-label similarity. High analytical flexibility therefore contributes to the observed lexical dispersion in how researchers describe these steps.

This analysis provides an internal assessment of the annotation schema. The high within-label similarity of standardized preprocessing labels demonstrates that these classes group text snippets with consistent methodological content. The lower similarity of graph-analysis labels identifies areas of the schema in which lexical matching alone would be insufficient for reliable extraction. These categories require contextual interpretation because the same analytical decision may be expressed through different terminology, narrative structures, and parameter descriptions. This linguistic variability demonstrates why automated extraction requires advanced contextual models rather than basic keyword matching.

The lexical-consistency results also help explain why functional-connectivity and graph-analysis annotations showed the greatest improvement after prompt refinement. Explicit definitions and schema constraints were particularly important for labels whose relevant information could not be identified reliably through recurring keywords alone.

### Overall validation outcome

The two validation components address different dimensions of dataset quality. Alignment with the manually curated extraction resource showed that the automated workflow recovered a substantial proportion of previously identified methodological information and that prompt refinement markedly improved annotation completeness, particularly for functional-connectivity and graph-analysis decisions. The lexical-consistency analysis showed that established preprocessing labels formed coherent annotation families, while also identifying graph-analysis labels for which lower lexical similarity reflected genuine variation in methodological description.

All discrepancies identified through automated structural checks, comparison with the manual extraction resource, prompt evaluation, and within-label consistency analysis were examined during the final expert validation round. Corrections were made with reference to the original article text, after which the verified annotations were exported in structured JSON format.

Together, these procedures establish the external agreement, internal coherence, and source-level traceability of the ASTEROID dataset. The validation framework assesses not only whether the automated annotations align with an existing manual extraction, but also whether the schema groups comparable methodological content across the corpus and appropriately represents areas in which reporting is intrinsically heterogeneous. The final dataset therefore provides a validated structured representation of analytical decisions reported in graph theory-based fMRI studies and a reference resource for the evaluation and development of future methodological-information extraction systems.

## Usage Notes

The following notes provide practical guidance for reusing and interpreting the dataset. They are intended to clarify the scope of the dataset, the role of the annotation schema, and the key limitations of reporting practices in the source literature.

### Scope of reuse

This dataset provides structured annotations of analytical decision descriptions reported in graph theory–based fMRI studies, together with study-characteristics information for interpreting the source articles. Users should reuse the dataset within this domain-specific scope and avoid generalizing the annotations to neuroimaging studies more broadly without additional validation.

### Interpretation of analytical decisions

Users are advised to consult the schema documentation before interpreting individual step definitions, software entries, algorithm/type labels, and parameter fields. Because methodological reporting in the source literature is often incomplete, narrative, or implicit, the absence of an annotated item should not automatically be interpreted as evidence that a step was not performed. It may instead reflect insufficient or ambiguous reporting in the original article.

### Use of study-characteristics information

The study-characteristics labels are provided to support interpretation of the annotated analytical decisions. These labels include information such as publication details, participant characteristics, acquisition parameters, and task or experimental design features. They should be treated as study-characteristics descriptors rather than as the primary focus of the dataset.

### Use for automated extraction methods

The dataset may be used as a structured reference for testing or comparing automated information-extraction methods within the domain of graph theory–based fMRI research. Users should account for the dataset size, annotation schema, and domain specificity when interpreting such evaluations.

### Versioning and documentation

Users should refer to the repository documentation, schema files, and version history when reusing the dataset. Any updates to the dataset or scripts should be cited according to the version used, once persistent identifiers are available.

## Data Availability

The ASTEROID dataset has been deposited as a draft on Zenodo under the reserved DOI (https://doi.org/10.5281/zenodo.23569927) The dataset is currently under review and is not yet publicly accessible. Upon completion of the necessary release requirements, the dataset and accompanying documentation will be made publicly available through Zenodo and GitHub (https://github.com/sanahassanimam/ASTEROID-Validated-Annotated-Dataset). The repository will include the structured annotation files, schema documentation, processing scripts, and supplementary materials required for reuse and interpretation.

## Code Availability

All custom scripts used for annotation merging, quality control, mismatch analysis, and review support will be made publicly available through the ASTEROID GitHub repository (https://github.com/sanahassanimam/ASTEROID-Validated-Annotated-Dataset) upon dataset release.

The PDF-to-JSON conversion script is already publicly available in the separate Annotation-buddy GitHub repository (https://github.com/sanahassanimam/Annotation-buddy).

Supporting scripts are provided in the Python_Scripts/ directory. These scripts document and support the main processing steps used to prepare and validate the final dataset:

**mismatch_checker.py**: performs mismatch analysis between the structured JSON annotations and the manually curated reference information. It identifies matched records, non-matched records, entries present only in the manual reference, and entries present only in the JSON annotations.

**combine_summary.py:** generates summary outputs from the annotation and validation results. These summaries support inspection of extracted information across studies and help identify patterns, missing values, or inconsistencies.

**merge_json_file.py:** merges JSON files generated from the first extraction prompt and the graph-analysis-specific prompt into a combined output file. This step was used to integrate graph-analysis information that was missing or incomplete in the first-prompt output, thereby supporting comparison and consolidation across prompt versions.

**review_newversion.py:** supports the review process by organizing validation outputs and highlighting entries that require human inspection. This script was used to guide the expert review and correction of annotations before the final dataset release.

The scripts were developed and tested between August 2025 and December 2025 using Python 3.10. Key dependencies include pandas, numpy, json, pathlib, re, and visualization packages used for generating validation summaries and manuscript figures. A requirements.txt file containing the complete software environment and package versions will be provided in the repository.

The repository will be released under an appropriate open license, with code and data/documentation licenses specified separately where required. The released scripts reproduce the annotation merging, mismatch analysis, quality-control summaries, and validation outputs reported in the manuscript.

Together, these scripts and structured output folders support transparency and reproducibility of the dataset preparation process by documenting how annotations were generated, compared, merged, reviewed, and prepared for final validation.

## Author Contributions

SI: conceptualization, data curation, methodology, software, formal analysis, visualization, writing – original draft, DC: data curation, methodology, software, formal analysis, DK, JW & CS: data curation, methodology, visualization, writing – review and editing, AH: conceptualization, methodology, supervision, writing – review and editing, funding acquisition, CG: conceptualization, data curation, methodology, supervision, writing – review and editing, funding acquisition.

